## Supplementary material for "Affinity-tagged SMAD1 and SMAD5 mouse lines reveal transcriptional reprogramming mechanisms during early pregnancy": Revision2_SuppFile1

### BETA: MOTIF ANALYSIS

Motif Scan on the TF Target Genes

#### PART1: UP TARGET GENES

| Symbol | DNA BindDom | Species | Pvalue (T Test) | T Score | Logo |
|---|---|---|---|---|---|
| Myb<br>Mybl1 | Myb Domain Family<br>Myb Domain Family | Mus musculus | 1.85e-02 | 2.09 | 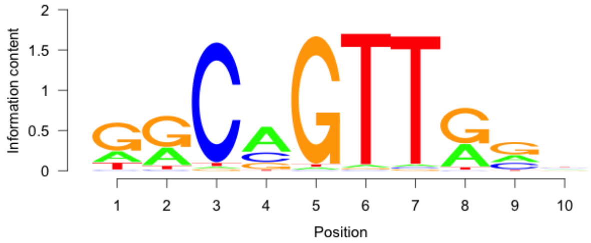 |
| Hmbox1 | Homeodomain Family | Mus musculus | 2.85e-02 | 1.90 | 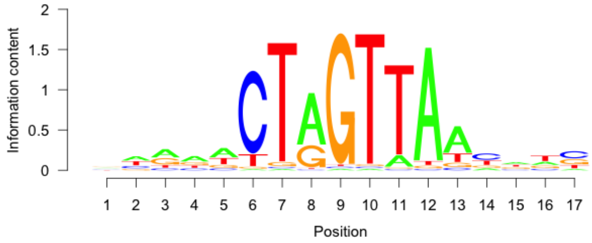 |
| Klf4<br>Klf1<br>Klf12<br>Klf7 | BetaBetaAlpha-zinc finger Family<br>BetaBetaAlpha-zinc finger Family<br>BetaBetaAlpha-zinc finger Family<br>BetaBetaAlpha-zinc finger Family | Mus musculus | 3.75e-02 | 1.78 | 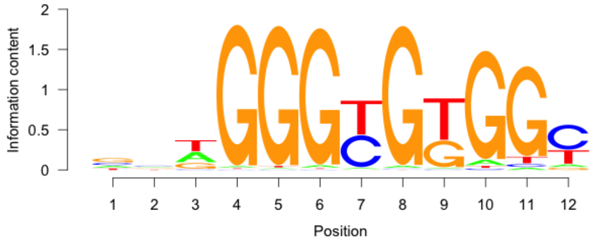 |
| Gsc | Homeodomain Family | Mus musculus | 4.34e-02 | 1.71 | 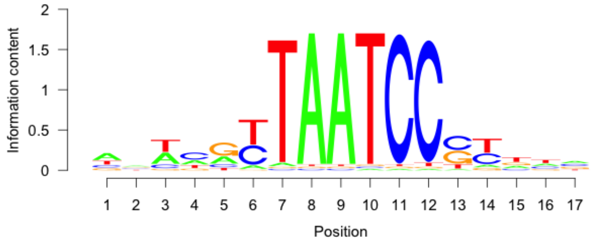 |
| Sox1 | High Mobility Group (Box) Family | Mus musculus | 6.75e-02 | 1.50 | 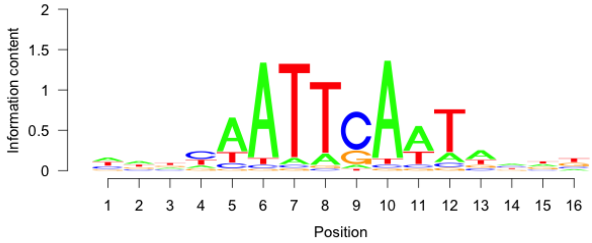 |
| Rhox11 | Homeodomain Family | Mus musculus | 8.37e-02 | 1.38 | |

### PART3: UP VS DOWN MOTIF SCAN

| Symbol | DNA BindDom | Species | Pvalue (T Test) | T Score | Logo |
|---|---|---|---|---|---|
| Uncx<br>Nkx6-1<br>Hoxd9 | Homeodomain Family<br>Homeodomain Family<br>Homeodomain Family | Mus musculus | 1.05e-04 | 3.89 | 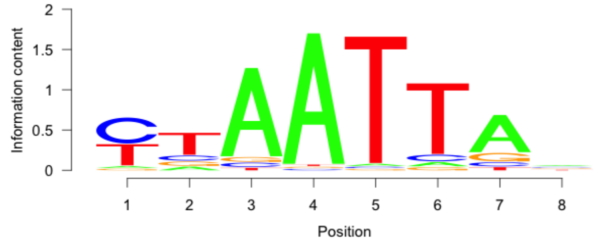 |
| Zfp423 | BetaBetaAlpha-zinc finger Family | Mus musculus | 6.57e-04 | -3.41 | 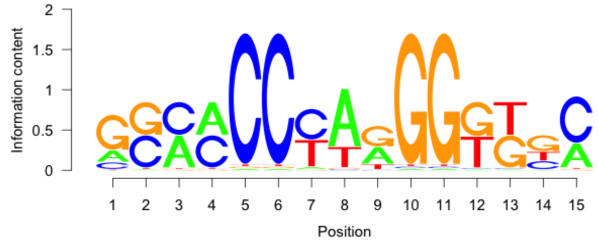 |
| Etv3<br>Elk3 | Ets Domain Family<br>Ets Domain Family | Mus musculus | 2.02e-03 | 3.09 | 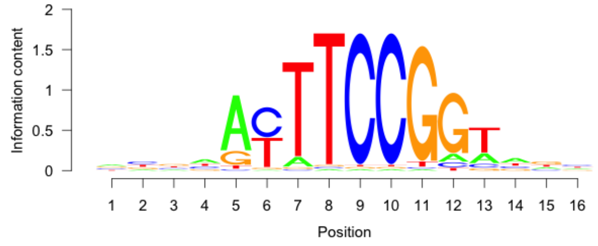 |
| Hoxd8<br>Sox17 | Homeodomain Family<br>High Mobility Group (Box) Family | Mus musculus | 2.43e-03 | -3.04 | 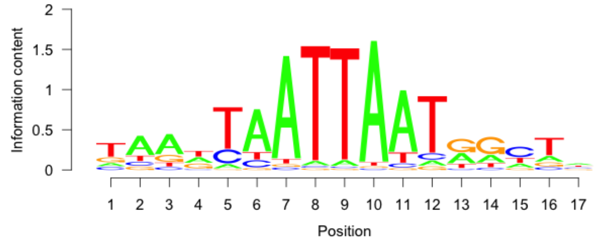 |
| Creb5 | Leucine Zipper Family | Mus musculus | 4.47e-03 | 2.85 | 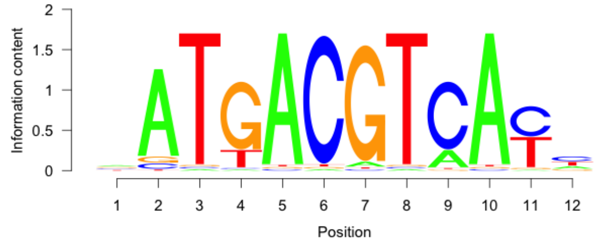 |
| Rfx2 | RFX Domain Family | Mus musculus | 5.98e-03 | 2.75 | 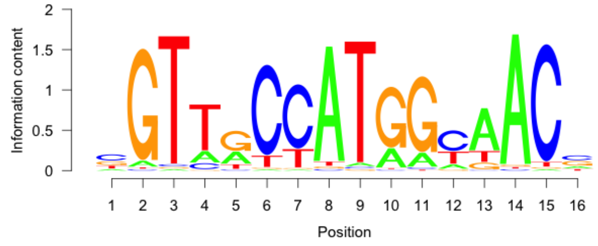 |

| | | | | | |
|---|---|---|---|---|---|
| Hoxb4 | Homeodomain Family | Mus musculus | 8.08e-02 | 1.75 | 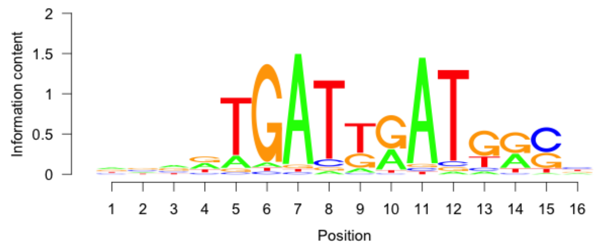 |
| Rfx2 | RFX Domain Family | Mus musculus | 1.20e-01 | -1.55 | 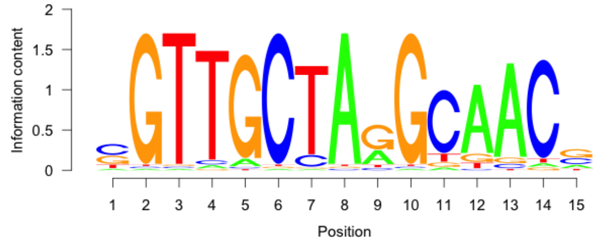 |
| Lhx8 | Homeodomain Family | Mus musculus | 1.20e-01 | -1.55 | 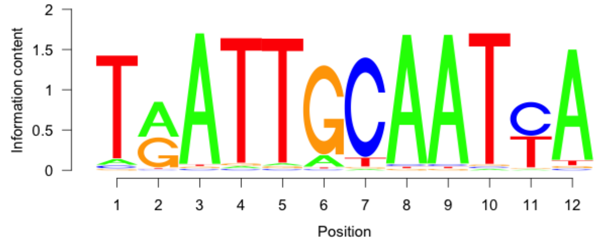 |

#### PART4: UP VS NON TARGET MOTIF

| Symbol | DNA BindDom | Species | Pvalue (T Test) | T Score | Logo |
|---|---|---|---|---|---|
| Uncx | Homeodomain Family | Mus musculus | 3.96e-09 | 5.94 | 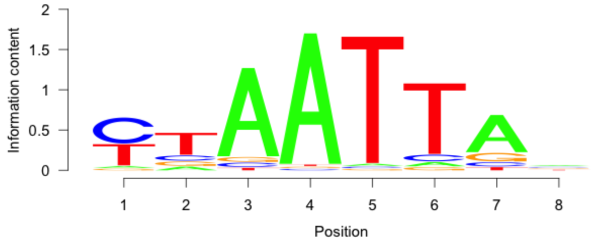 |
| Etv3<br>Ets1 | Ets Domain Family<br>Ets Domain Family | Mus musculus | 6.53e-06 | 4.53 | 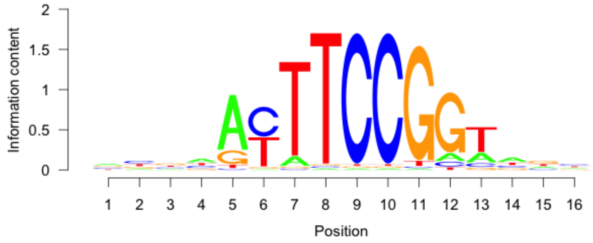 |
| Olig2 | Helix-Loop-Helix Family | Mus musculus | 9.17e-06 | 4.46 | 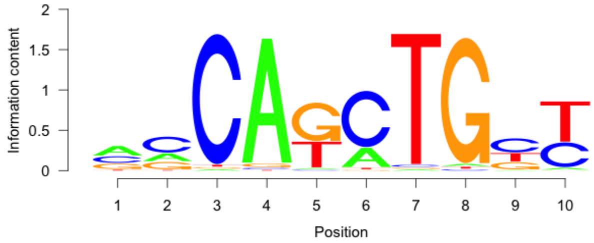 |

| | | | | | |
|---|---|---|---|---|---|
| Olig2 | Helix-Loop-Helix Family | Mus musculus | 3.07e-04 | 3.62 | 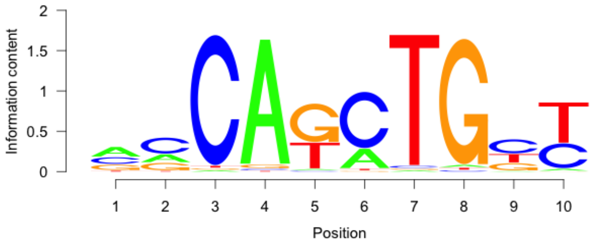 |
| Batf | Leucine zipper Family | Mus musculus | 5.92e-04 | 3.44 | 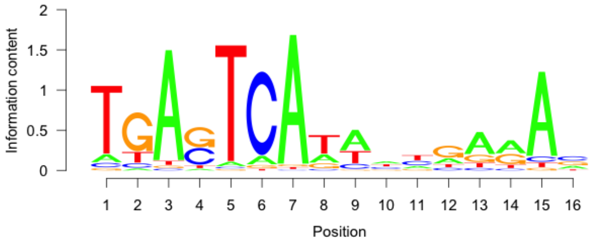 |
| Otx1 | Homeodomain Family | Mus musculus | 2.77e-03 | 3.00 | 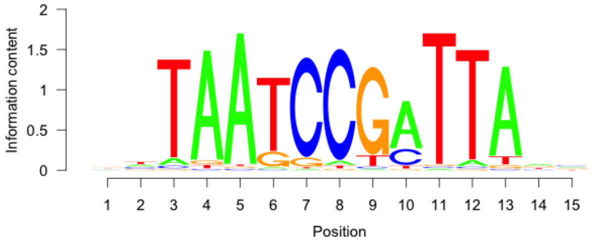 |
| Nanog | Homeodomain Family | Mus musculus | 3.01e-03 | 2.97 | 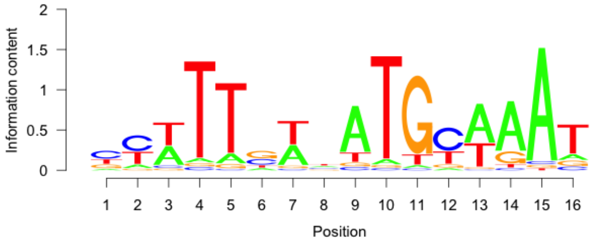 |
| Nkx3-1 | Homeodomain Family | Mus musculus | 3.05e-03 | 2.97 | 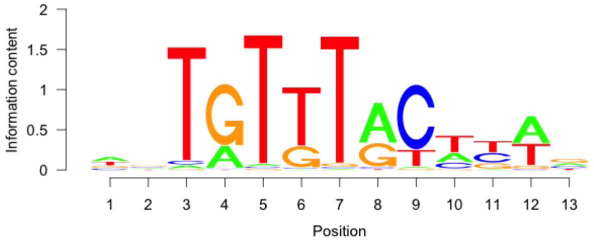 |
| Hoxd3 | Homeodomain Family | Mus musculus | 4.23e-03 | 2.87 | 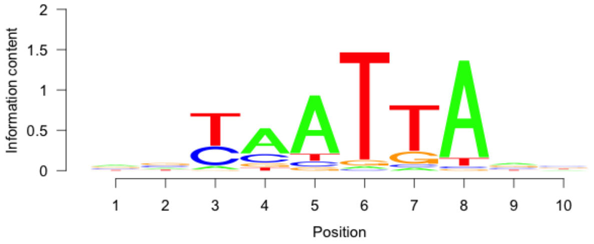 |
| Sox3 | High Mobility Group (Box) Family | Mus musculus | 5.25e-03 | 2.80 | 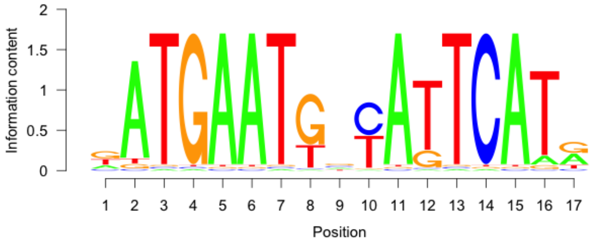 |
