## Supplementary material for "Affinity-tagged SMAD1 and SMAD5 mouse lines reveal transcriptional reprogramming mechanisms during early pregnancy": Revision2_SuppFile2

### BETA: MOTIF ANALYSIS

Motif Scan on the TF Target Genes

#### PART1: UP TARGET GENES

| Symbol | DNA BindDom | Species | Pvalue (T Test) | T Score | Logo |
|---|---|---|---|---|---|
| Ebf1 | Helix-Loop-Helix Family | Mus musculus | 1.57e-02 | 2.15 | 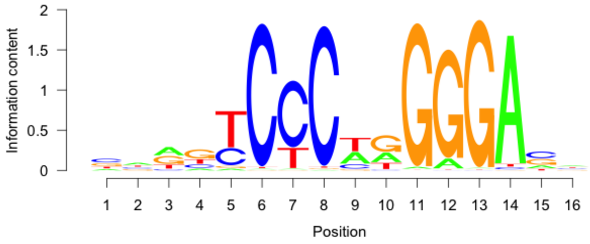 |
| Zfp128 | BetaBetaAlpha-zinc finger Family | Mus musculus | 5.24e-02 | 1.62 | 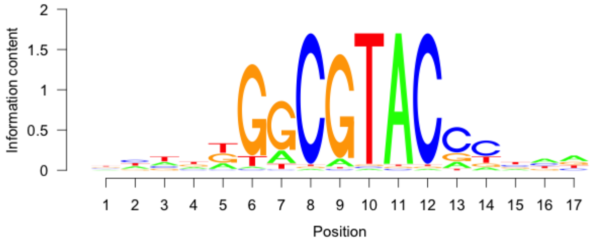 |
| Otx1 | Homeodomain Family | Mus musculus | 5.38e-02 | 1.61 | 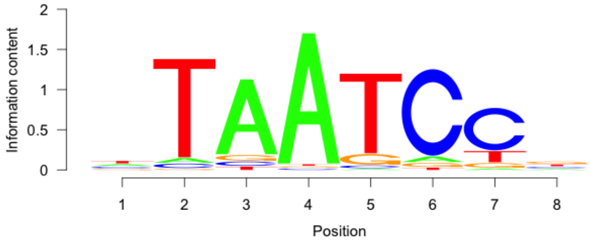 |
| Rela | Rel Homology Region Family | Mus musculus | 5.53e-02 | 1.60 | 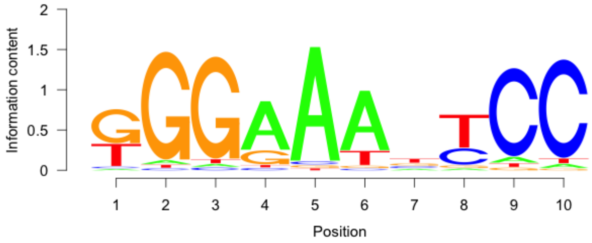 |
| Tp53 | Loop-Sheet-Helix Family | Mus musculus | 5.87e-02 | 1.57 | 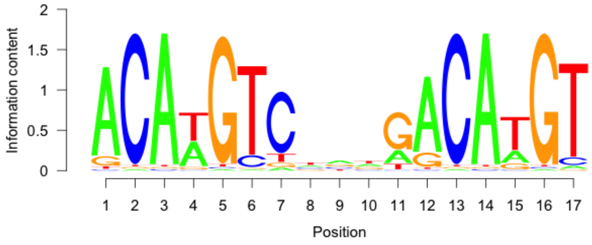 |
| Tcfap2e<br>Tcfap2c | Helix-Loop-Helix Family<br>Helix-Loop-Helix Family | Mus musculus | 6.92e-02 | 1.48 | |

| | | | | | |
|---|---|---|---|---|---|
| | | | | | 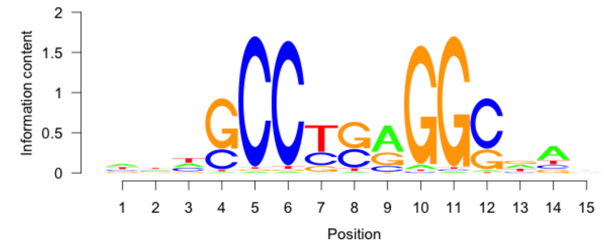 |
| Nr2e1 | Hormone-nuclear Receptor Family | Mus musculus | 8.68e-02 | 1.36 | 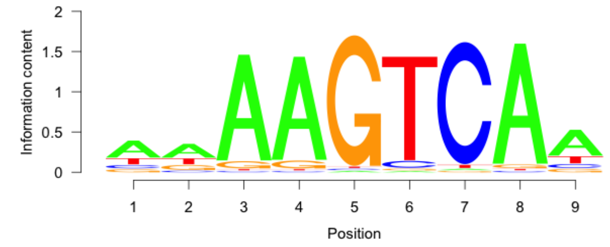 |
| Srf | MADS Box Family | Mus musculus | 1.16e-01 | 1.19 | 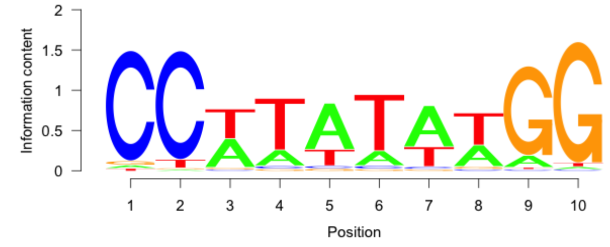 |
| Mybl1 | Myb Domain Family | Mus musculus | 1.50e-01 | 1.04 | 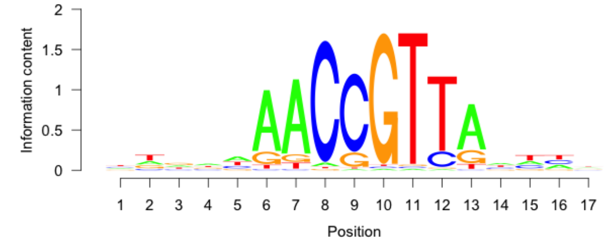 |

#### PART2: DOWN TARGET GENES

| Symbol | DNA BindDom | Species | Pvalue (T Test) | T Score | Logo |
|---|---|---|---|---|---|
| Arx<br>Isx<br>Pax7<br>Lbx2<br>Lhx9<br>Lhx1 | Homeodomain Family<br>Homeodomain Family<br>Homeodomain Family<br>Homeodomain Family<br>Homeodomain Family<br>Homeodomain Family | Mus musculus | 2.60e-02 | 1.94 | 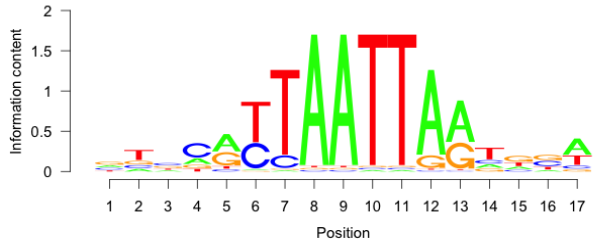 |
| Egr3 | BetaBetaAlpha-zinc finger Family | Mus musculus | 4.43e-02 | 1.70 | 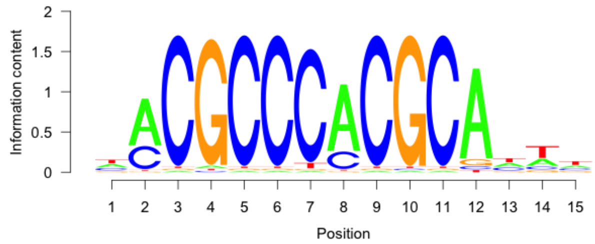 |

| | | | | | |
|---|---|---|---|---|---|
| Pax5 | Homeodomain Family | Mus musculus | 4.65e-02 | 1.68 | 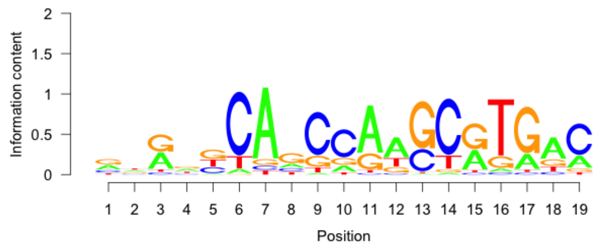 |
| Hoxd10 | Homeodomain Family | Mus musculus | 6.83e-02 | 1.49 | 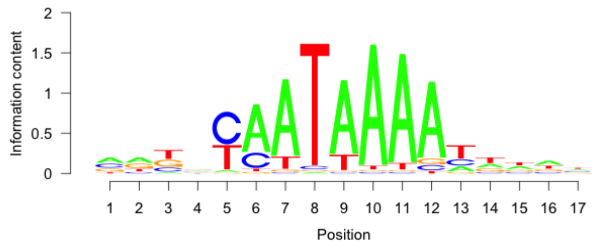 |
| Hoxd11 | Homeodomain Family | Mus musculus | 8.41e-02 | 1.38 | 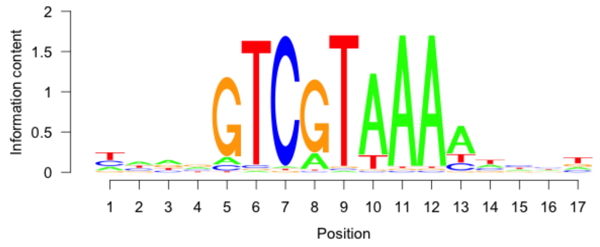 |

#### PART3: UP VS DOWN MOTIF SCAN

| Symbol | DNA BindDom | Species | Pvalue (T Test) | T Score | Logo |
|---|---|---|---|---|---|
| Creb3l2<br>Jdp2 | Leucine Zipper Family<br>Leucine Zipper Family | Mus musculus | 3.58e-02 | 2.10 | 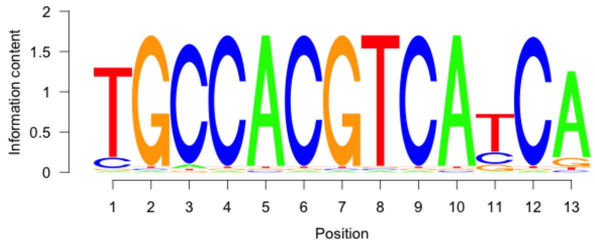 |
| Spi1 | Ets Domain Family | Mus musculus | 6.40e-02 | -1.85 | 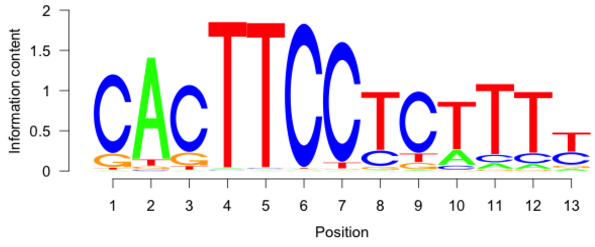 |
| Nkx3-1<br>Gbx1 | Homeodomain Family<br>Homeodomain Family | Mus musculus | 6.44e-02 | -1.85 | 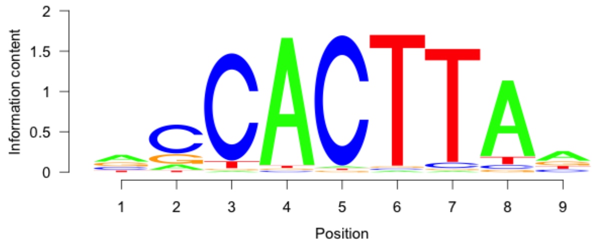 |

| | | | | | |
|---|---|---|---|---|---|
| Foxj3 | Forkhead Domain Family | Mus musculus | 7.96e-02 | -1.75 | 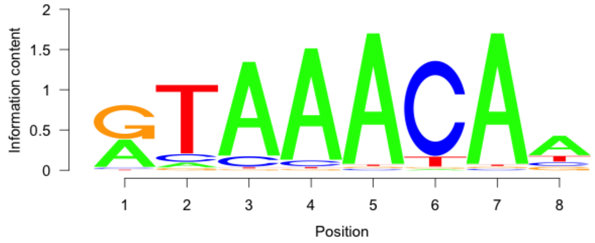 |
| Nr2e1 | Hormone-nuclear Receptor Family | Mus musculus | 9.92e-02 | -1.65 | 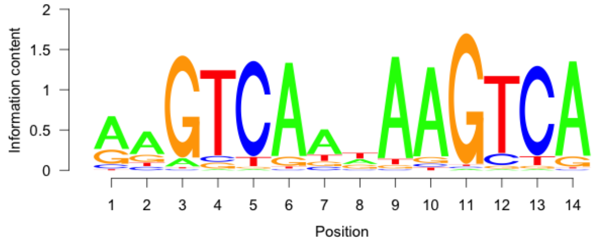 |
| Foxj3<br>Foxg1 | Forkhead Domain Family<br>Forkhead Domain Family | Mus musculus | 1.05e-01 | 1.62 | 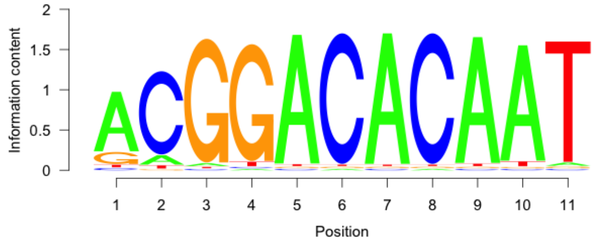 |
| Mybl1 | Myb Domain Family | Mus musculus | 1.05e-01 | 1.62 | 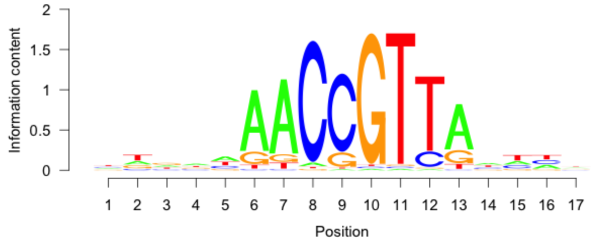 |
| Hmbox1 | Homeodomain Family | Mus musculus | 1.14e-01 | 1.58 | 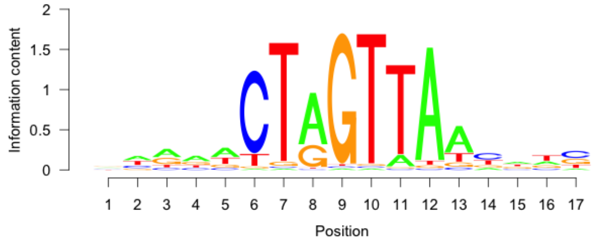 |
| Elk3<br>Gm5454 | Ets Domain Family<br>Ets Domain Family | Mus musculus | 1.33e-01 | 1.50 | 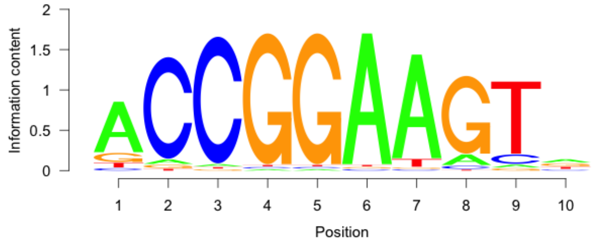 |
| Pax7<br>Hoxd8 | Homeodomain Family<br>Homeodomain Family | Mus musculus | 1.72e-01 | -1.37 | 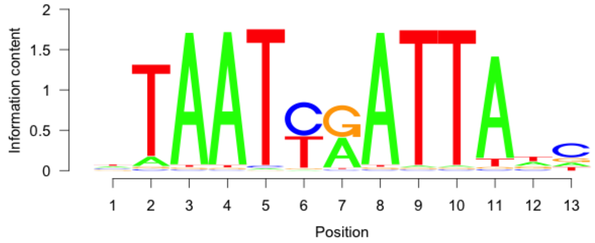 |

| | | | | | |
|---|---|---|---|---|---|
| Max | Helix-Loop-Helix Family | Mus musculus | 1.78e-01 | -1.35 | 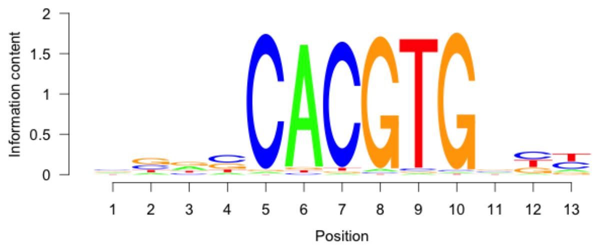 |
| Irf4 | Interferon Regulatory Factor | Mus musculus | 1.87e-01 | 1.32 | 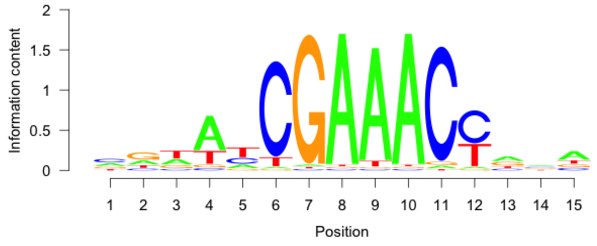 |
| Jun | Leucine Zipper Family | Mus musculus | 1.87e-01 | -1.32 | 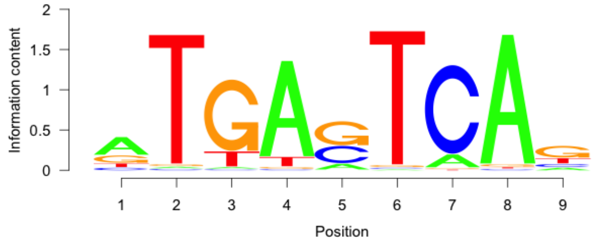 |
| Pknox2 | Homeodomain Family | Mus musculus | 1.94e-01 | -1.30 | 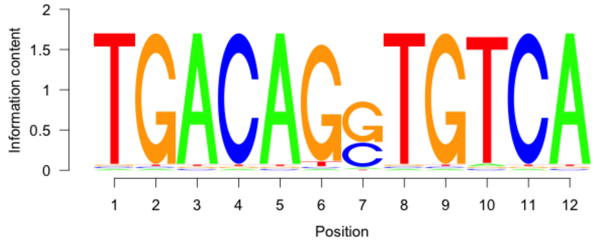 |
| Nr2f6 | Hormone-nuclear Receptor Family | Mus musculus | 2.11e-01 | 1.25 | 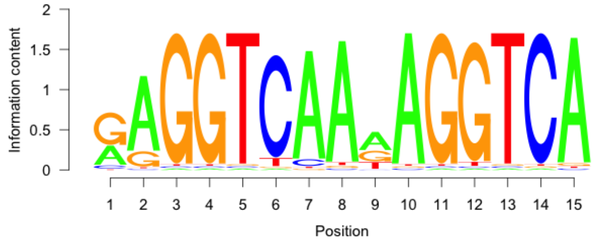 |

#### PART4: UP VS NON TARGET MOTIF

| Symbol | DNA BindDom | Species | Pvalue (T Test) | T Score | Logo |
|---|---|---|---|---|---|
| Rfx2 | RFX Domain Family | Mus musculus | 1.18e-07 | 5.32 | 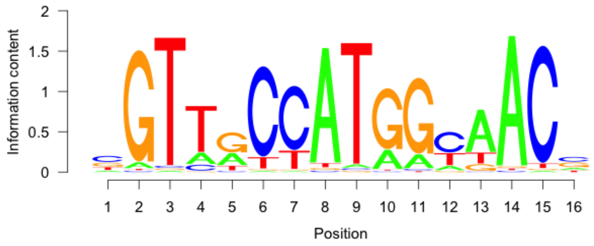 |

| | | | | | |
|---|---|---|---|---|---|
| Etv3<br>Elk3 | Ets Domain Family<br>Ets Domain Family | Mus musculus | 3.40e-05 | 4.16 | 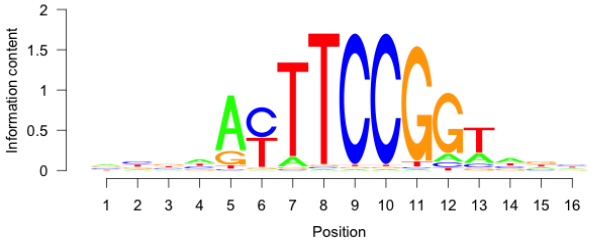 |
| Tal1 | Helix-Loop-Helix Family | Mus musculus | 1.09e-04 | 3.88 | 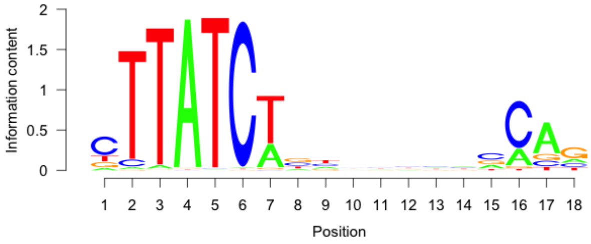 |
| Ctcf | BetaBetaAlpha-zinc finger Family | Mus musculus | 1.65e-03 | 3.15 | 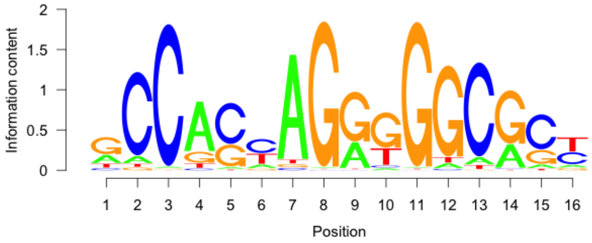 |
| Dbp<br>Jdp2 | Leucine Zipper Family<br>Leucine Zipper Family | Mus musculus | 5.06e-03 | 2.81 | 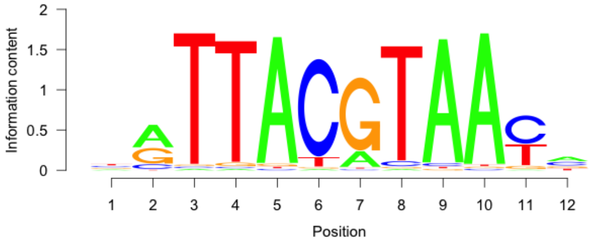 |
| Nanog | Homeodomain Family | Mus musculus | 1.10e-02 | 2.54 | 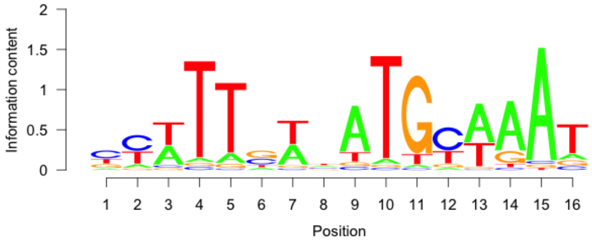 |
| Foxg1 | Forkhead Domain Family | Mus musculus | 1.33e-02 | 2.48 | 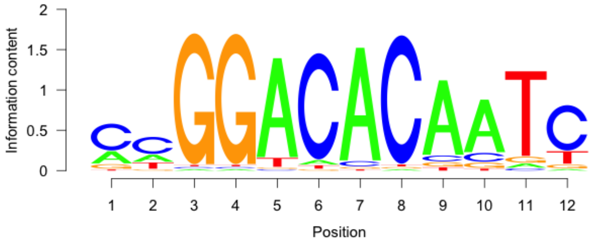 |
| Rela | Rel Homology Region Family | Mus musculus | 2.17e-02 | 2.30 | 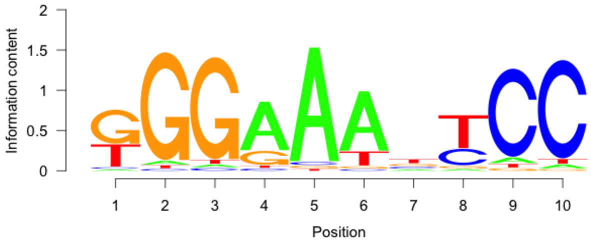 |

#### PART5: DOWN VS NON TARGET MOTIF

| Symbol | DNA BindDom | Species | Pvalue (T Test) | T Score | Logo |
|---|---|---|---|---|---|
| Etv3 | Ets Domain Family | Mus musculus | 2.57e-08 | 5.60 | 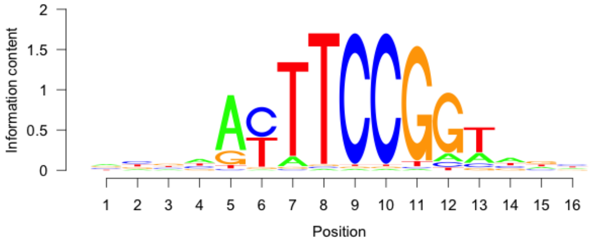  |
| Rfx2 | RFX Domain Family | Mus musculus | 1.42e-06 | 4.84 | 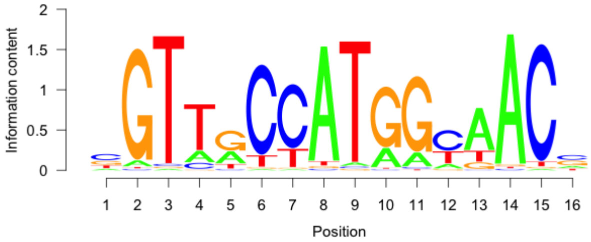  |
| Tal1 | Helix-Loop-Helix Family | Mus musculus | 1.04e-03 | 3.28 | 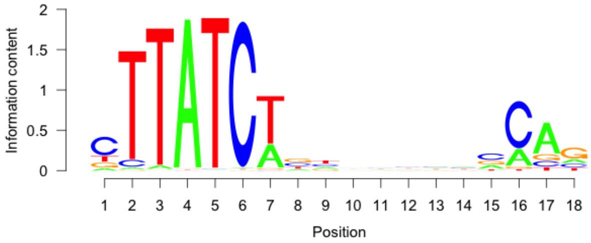  |
| Olig2 | Helix-Loop-Helix Family | Mus musculus | 3.85e-03 | 2.89 | 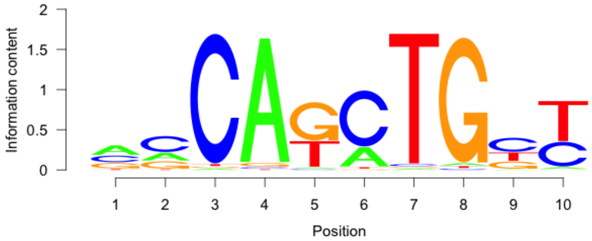  |
| Osr2 | BetaBetaAlpha-zinc finger Family | Mus musculus | 4.15e-03 | 2.87 | 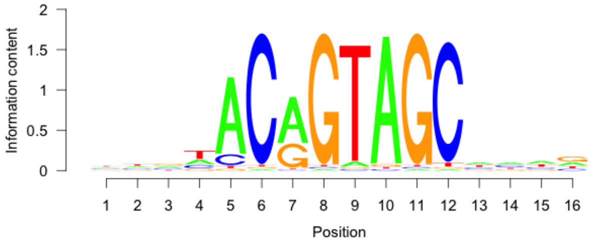  |
| Otx1 | Homeodomain Family | Mus musculus | 1.10e-02 | 2.54 | 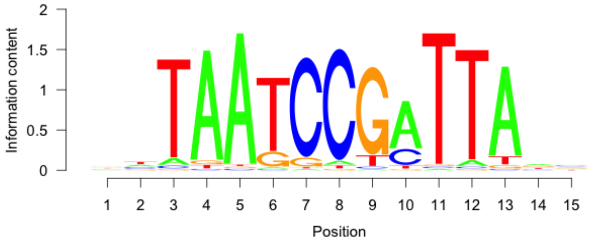  |

| | | | | | |
|---|---|---|---|---|---|
| Ctcf | BetaBetaAlpha-zinc finger Family | Mus musculus | 1.61e-02 | 2.41 | 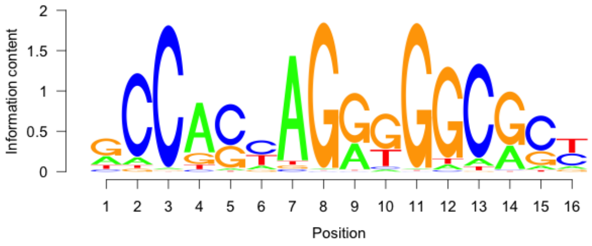  |
| Rela | Rel Homology Region Family | Mus musculus | 3.40e-02 | 2.12 | 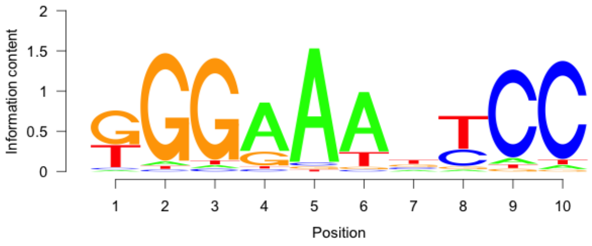  |
| Uncx | Homeodomain Family | Mus musculus | 3.98e-02 | 2.06 | 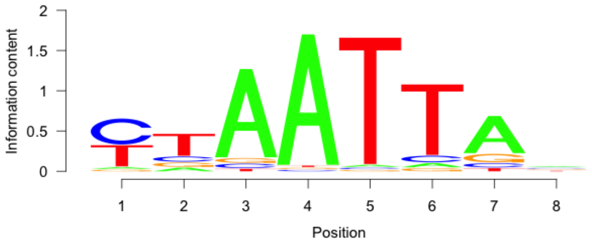  |
| Dbp | Leucine Zipper Family | Mus musculus | 4.14e-02 | 2.04 | 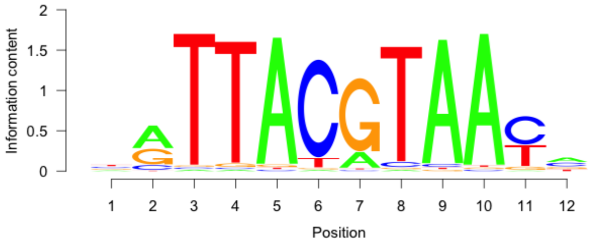  |
